# Trait Evolution with Jumps: Illusionary Normality

**DOI:** 10.1101/188854

**Authors:** Krzysztof Bartoszek

## Abstract

Phylogenetic comparative methods for real-valued traits usually make use of stochastic process whose trajectories are continuous. This is despite biological intuition that evolution is rather punctuated than gradual. On the other hand, there has been a number of recent proposals of evolutionary models with jump components. However, as we are only beginning to understand the behaviour of branching Ornstein–Uhlenbeck (OU) processes the asymptotics of branching OU processes with jumps is an even greater unknown. In this work we build up on a previous study concerning OU with jumps evolution on a pure birth tree. We introduce an extinction component and explore via simulations, its effects on the weak convergence of such a process. We furthermore, also use this work to illustrate the simulation and graphic generation possibilities of the mvSLOUCH package.

## INTRODUCTION

Contemporary stochastic differential equation (SDE) models of continuous trait evolution are focused around the Ornstein-Uhlenbeck process [15]

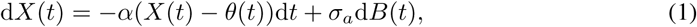

where *θ*(*t*) can be piecewise linear. These models take into account the phylogenetic structure between the contemporary species. The trait follows the SDE along each branch of the tree (with possibly branch specific parameters). At speciation times this process divides into two processes evolving independently from that point.

However, the fossil record indicates [10,13,14] that change is not as gradual as Eq. (1) suggests. Rather, that jumps occur and that the framework of Lévy processes could be more appropriate. There has been some work in the direction of phylogenetic Laplace motion [2,9,16] and jumps at speciation points [3,4,7,8]. In this work we describe recent asymptotic results on such a jump model (developed in [4]) and develop them by including an extinction component.

It is worth pointing out that OU with jumps (OUj) models are actually very attractive from a biological point of view. They seem to capture a key idea behind the theory of punctuated equilibrium (i.e. the theory of evolution with jumps [13]). At a branching event something dramatic (the jump) could have occurred that drove species apart. But then “The further removed in time a species from the original speciation event that originated it, the more its genotype will have become stabilized and the more it is likely to resist change.” [17]. Therefore, between branching events (and jumps) we could expect stasis—“fluctuations of little or no accumulated consequence” taking place [14]. This fits well with an OUj model. If ***α*** is large enough, then the process approaches its stationary distribution rapidly and the stationary oscillations around the (constant) mean can be interpreted as stasis between jumps.

In applications the phylogeny is given (from molecular sequences) but when the aim is to study large sample properties some model of growth has to be assumed. A typical one is the constant rate birth-death process conditioned on *n* contemporary tips (cf. [11,12,19]).

Regarding the phenotype the Yule-Ornstein-Uhlenbeck with jumps (YOUj) model was recently considered [4]. The phylogeny is a pure birth process (no extinction) and the trait follows an OU process. However, just after the *k*-th branching point (*k* = 1,…, *n* − 1, counting from the root with the root being the first branching point) on the phylogeny, with a probability *p*_*k*_, independently on each daughter lineage, a jump can occur. The jump is assumed to be normally distributed with mean 0 and variance 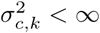. Just after a speciation event at time *t*, independently for each daughter lineage, the trait value *X*(*t*) will be

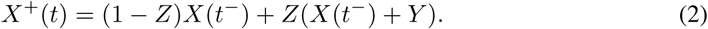

In the above Eq. (2) *X*(*t*^−/+^) means the value of *X*(*t*) respectively just before and after time *t, Z* is a binary random variable with probability *p*_*k*_ of being 1 (i.e. jump occurs) and 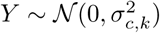.

## THE PURE BIRTH CASE

In the case of the pure birth tree central limit theorems (CLTs) can be explicitly derived thanks to a key property of the pure birth process. The time between speciation events *k* and *k* + 1 is exponential with parameter *λk* due to the memoryless property of the process and the law of the minimum of i.i.d. exponential random variables. This allows us to calculate Laplace transforms of relevant speciation times and count speciation events on lineages [3,4,6,19].

We first remind the reader about the mathematical concept of sequence convergence with density 1 (see e.g. [18]) and then summarize the previously obtained CLTs.

### Definition 1.

*A subset E* ⊂ ℕ *of positive integers is said to have density* 0 *if*

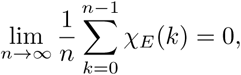

*where χ*_*E*_(·) *is the indicator function of the set E.*

### Definition 2.

*A sequence a*_*n*_ *converges to* 0 *with density* 1 *if there exists a subset E* ⊂ ℕ *of density* 0 *such that*

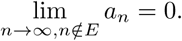

Let 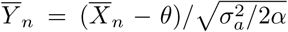 be the normalized sample mean of the YOUj process with 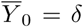. Denote by *y*_*n*_ the σ-algebra containing information on the tree and jump pattern (*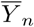* conditional on *y*_*n*_ is normal). Assume also that λ = 1. The restriction of λ = 1 is a mild one, changing λ is equivalent to rescaling time.

### Theorem 1 ([4]).

*Assume that the jump probabilities and jump variances are constant equalling *p* and 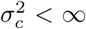 respectively. The process 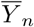 has the following, depending on* α, *asymptotic with n behaviour.*

I. *If* 0.5 *< α, then the conditional variance of the scaled sample mean 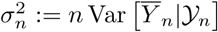 converges in* ℙ *to a random variable 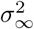 with mean*

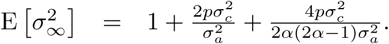 *The scaled sample mean, 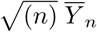 converges weakly to a random variable whose characteristic function can be expressed in terms of the Laplace transform of 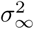*

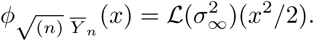
II. *If* 0.5 = *α, then the conditional variance of the scaled sample mean 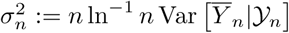 converges in* ℙ *to a random variable 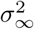 with mean*

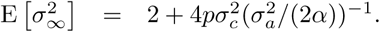 *The scaled sample mean, 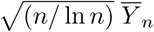 converges weakly to a random variable whose characteristic function can be expressed in terms of the Laplace transform of 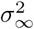*

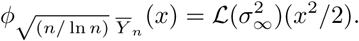
III. *If* 0 *< α <* 0.5, *then 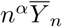 converges almost surely and in L^2^ to a random variable Y_α,δ_ with first two moments*

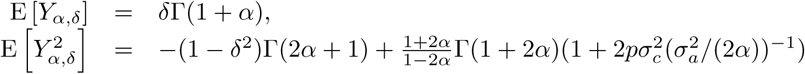

### Theorem 2 ([4]).

*If 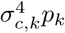 is bounded and goes to* 0 *with density* 1, *then depending on α the process 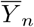 has the following asymptotic with n behaviour.*

I. *If* 0.5 *< α, then 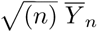 is asymptotically normally distributed with mean* 0 *and variance* (2*α* + 1)/(2*α* − 1).
II. *If* 0.5 = *α, then 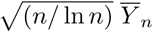 is asymptotically normally distributed with mean 0 and variance 2.*

**Remark 1.** *Notice that Thm. 2 immediately implies the CLTs when there are no jumps, i.e. p*_*k*_ = 0 *for all k* [6].

## EXTINCTION PRESENT

The no extinction assumption is difficult to defend biologically, unless one considers extremely young clades. Therefore, it is desirable to generalize Thms. 1 and 2 to the case when the extinction rate, *μ*, is non-zero. However, there are a number of intrinsic difficulties associated with such a generalization. We do not have the Laplace transforms of the time to coalescent of a random pair of tips (its expectation seems involved enough, [19]). More importantly we seem to be unable to say much about the number of hidden speciation events on a random lineage. We have to remember that a jump can be due to any speciation event, including those that lead to extinct lineages. A lineage that survived till today can have multiple branches stemming from it that faded away in the past, see Fig. 1.

**Figure 1.**
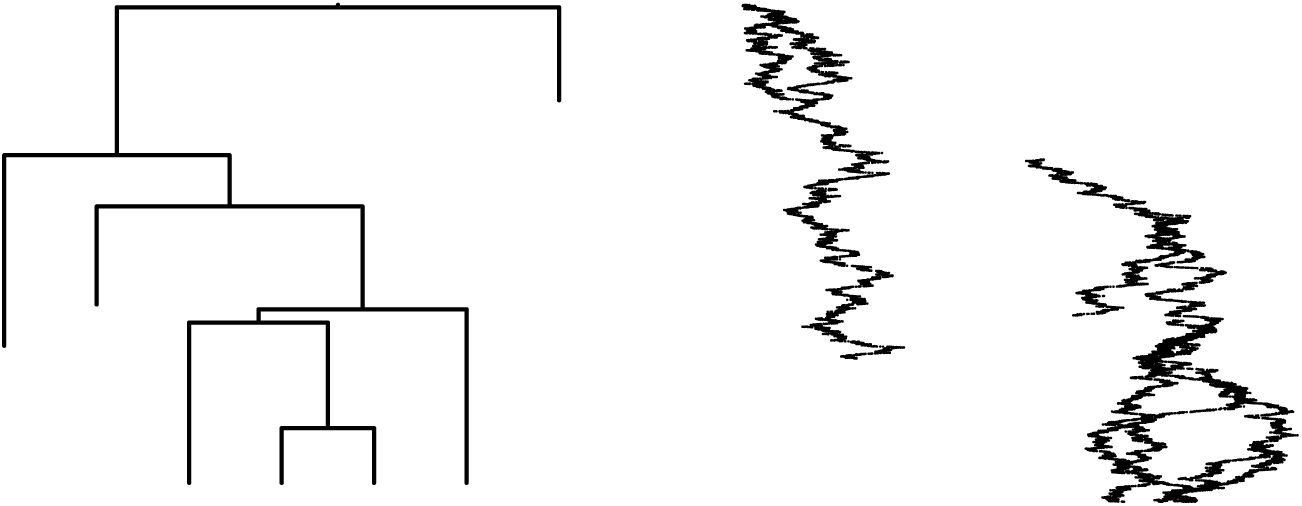
Left: birth-death tree with a clade of 4 contemporary species. Right: OUj process evolving on this tree (graphic by mvSLOUCH). There is a single jump in the trait process at the first speciation event after the root. The OU process is a slowly adapting one with large jump variance (*α* = 0.3, 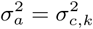 = 4, tree height: 2.427).

Furthermore, for the OU model of trait evolution, we need to know the distribution (or Laplace transform) of the times between the speciation events. Without extinction, *μ* = 0, this was simple. The time between speciation events *k* and *k* +1 was exponential with rate *λk*, as the minimum of *k* independent rate λ exponential random variables. However, when *μ* > 0 we not only need to know the number of hidden speciation events between two non-hidden (i.e. leading to contemporary tips) speciation events but also the law of the time between the hidden speciation. We do not know this law and furthermore as we are conditioning on *n* it is not entirely clear if times between speciation events will be independent (like they are in the pure birth case).

Our question is whether we can expect counterparts of Thms. 1 and 2 to hold when extinction is present, i.e. 0 < *μ* ≤ 1 = *λ*. We are not aware of analytical results on the issues raised in the previous paragraph. Hence, we will approach answering the problem by simulations. Based on the results in [1] we should expect a phase transition to occur for 1 − *μ* = *α*/2 (remember λ = 1).

We simulate 1000 birth-death trees for *μ* = 0.25,0.5,0.75 conditioned on *n* = 500 contemporary tip species with the TreeSim [20,21] R package. On each phylogeny we simulate an OU process with *α* = (1 − *μ*), (1 − *μ*)/2, (1 − *μ*)/4 using the mvSLOUCH R package [5]. In all simulations 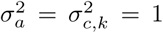, *p*_*k*_ = 0.5 (for all internal nodes) and *X*_0_ = *θ* = 0. For a given phylogeny, OU simulation pair we calculate the scaled sample average,

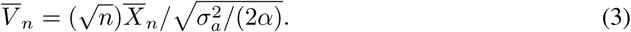

We report the results of the simulations by plotting histograms with a mean 0 and variance equalling the scaled sample variance normal curve.

## ILLUSIONARY NORMALITY?

We report the result of our simulations in Fig. 2 and Tab. 1. The histograms are not conclusive but they can be interpreted as indicating a similar as in Thms. 1 and 2 phase transition. In the fast adaptation case, *α* = (1 − *μ*) > (1 − *μ*)/2, the histograms do not seem to deviate much from the normal curve. On the other hand, it is a bit surprising that the “worst” looking histogram is when *μ* = 0.25, the one we would expect to be closest to “normality”. As the ratio of α to 1 − *μ* decreases we can see that the histograms of 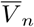 deviate more from the normal curve. Furthermore, when *α* = (1 − *μ*)/4 < (1 − *μ*)/2 we can start to see heavier than normal tails.

**Figure 2.**
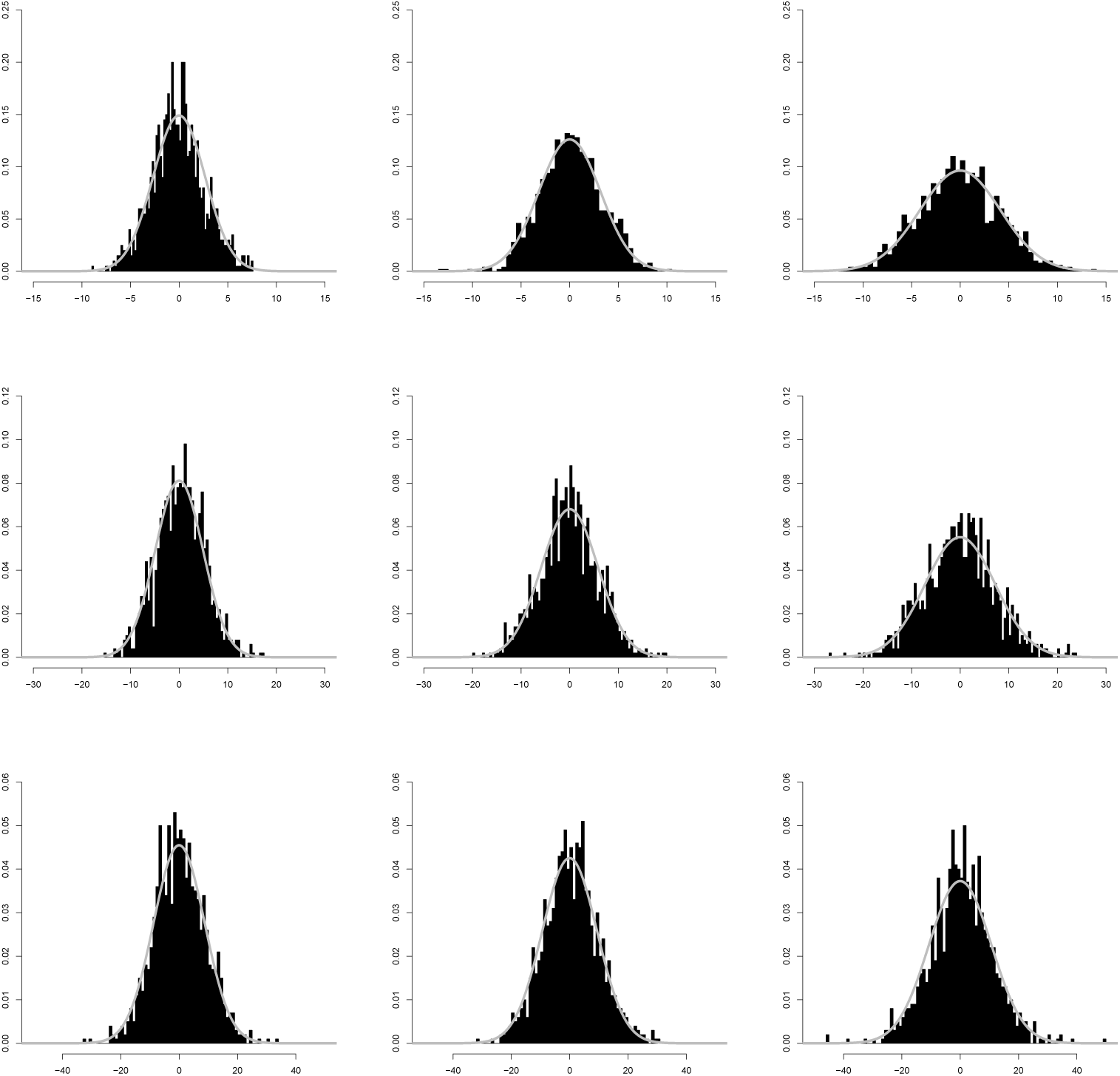
Simulations of Eq. (3). Left column *μ* = 0.25, centre column *μ* = 0.5, right column *μ* = 0.75, top row *α* = (1 − *μ*), centre row *α* = (1 − *μ*)/2 and bottom row α = (1 − *μ*)/4. The gray curve is the density curve of the normal distribution with mean 0 and variance equalling the sample variance. Each histogram is constructed from 1000 simulated birth-death trees (birth rate 1, death rate *μ*) with an OUj process (*θ = X*_0_ = 0, 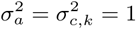, *p*_*k*_ =0.5) evolving on top of the tree. Notice that the *x* and *y* axes differ between rows.

**Table 1.** Summary of simulated samples. The mean, variance, skewness and excess kurtosis refer to the sample moments of 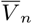. The 95% bootstrap confidence intervals for the excess kurtosis are based on 10000 bootstrap replicates (of the 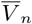 values) and calculated by the R package boot (basic bootstrap confidence intervals are reported).

| $\mu$ | $\alpha$ | $\alpha/(1 - \mu)$ | mean | variance | skewness | excess kurtosis |
| --- | --- | --- | --- | --- | --- | --- |
| 0.25 | 0.75 | 1 | −0.117 | 7.173 | 0.114 | 0.046 (−0.195, 0.27) |
| 0.5 | 0.5 | 1 | 0.116 | 10.016 | −0.084 | 0.280 (−0.297, 0.803) |
| 0.75 | 0.25 | 1 | −0.082 | 17.158 | 0.058 | −0.058 (−0.318, 0.186) |
| 0.25 | 0.375 | 0.5 | 0.344 | 24.227 | 0.038 | 0.110 (−0.185, 0.392) |
| 0.5 | 0.25 | 0.5 | −0.141 | 34.432 | 0.012 | 0.121 (−0.186, 0.412) |
| 0.75 | 0.125 | 0.5 | −0.056 | 52.357 | 0.087 | 0.3 (−0.064, 0.646) |
| 0.25 | 0.1875 | 0.25 | 0.212 | 77.016 | 0.072 | 0.316 (−0.107, 0.722) |
| 0.5 | 0.125 | 0.25 | 0.575 | 88.128 | 0.102 | 0.149 (−0.14, 0.416) |
| 0.75 | 0.0625 | 0.25 | 0.3 | 114.806 | −0.024 | 1.169 (0.328, 1.964) |

The analysis of the first four moments in Tab. 1 does point to two things. Firstly, the scaling by 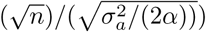 is incorrect (cf. Thm. 3.3 [1]). In all setups the sample variance of 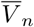 is much greater than 1. However, based on the estimates of skewness and excess kurtosis we cannot reject normality outright. Only in the most extreme case (*μ* = 0.75, *α*/(1 − *μ*) = 0.25) do the 95% bootstrap confidence intervals of the excess kurtosis not cover 0. When *α* = (1 − *μ*)/2 we should actually expect to have a logarithmic correction in the scaling, 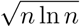, however we present here the histograms without it. Such a logarithmic correction did not bring in any qualitative changes to the results and hence, for easiness of comparison in Tab. 1 and Fig. 2 we refrain from using it.

Based on the histograms and analysis of the first four moments we would not suspect that with a constant jump probability *p* we do not have a nearly classical CLT, i.e. weak convergence to a normal limit after scaling by 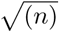. All that we would think would remain, would be to find the correct variance of the limit. In fact if we look at Fig. 2 in [4] we would be under a similar illusion. The histogram for *α* = 1, λ = 1, *μ* = 0, *p*_*k*_ =0.5 and 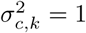 nearly perfectly fits into a normal density curve. However, Thms. 1 and 2 are very clear that for a constant product of 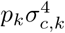 we will not have a normal limit. Therefore, with *μ* > 0 we cannot expect a change of the situation. On the one hand a non-zero extinction rate does cause the (conditional on *n* contemporary tips) tree to be higher. But on the other hand, with greater height comes to opportunity for more jumps. And this variability of jump occurrences seems to be the force pushing the limit away from normality.

The simulations and results presented here and in [4] should also serve as a warning. Visual inspection of histograms from simulated data is never sufficient for drawing conclusions about a model’s underlying distribution. A bare minimum are goodness-of-fit tests but their conclusions should be supported by analytical derivations.

## ACKNOWLEDGEMENTS

KB’s research was supported by the Knut and Alice Wallenberg Foundation. KB’s conference participation was supported by the Wenner-Gren Foundation (grant nr. RSh2016-0078).

